# Progesterone regulation of cervix ripening in preterm and term birth

**DOI:** 10.1101/2025.01.31.636012

**Authors:** Steven M. Yellon, Daylan Ward, Ashley Thompson, Brigitte M. Vazquez, D. Daniel Baldwin, Elaine J. Oldford, Michael A. Kirby

## Abstract

The cervix functions both as gatekeeper barrier to maintain pregnancy and virtually vanish for birth at term in mammals. The period of remodeling well-before term is characterized by an inflammatory process associated with reduced cell nuclei density and cross-linked collagen, as well as increased density of resident macrophages in cervix stroma. Contemporarily, progesterone (P4) is at or near peak concentrations in maternal circulation. The functional or actual loss of response to P4 is thought to drive the process that enhances uterine contractile activity for labor and parturition at term. The objective of the present study was to determine if actual or functional loss of P4 regulated cytomorphological characteristics associated with prepartum cervix ripening at term and with preterm birth. On day 16 of pregnancy. Ovaries were removed to eliminate the main source of P4 production and a silastic capsule implanted (with vehicle or P4, Ovx or Ovx+P4, respectively).

Controls received a vehicle-filled capsule, while a P4 capsule was implanted into an addition group of Intact mice to ensure sustained concentrations throughout pregnancy (Intact+P4). Pups were born in controls at term (days 19-20 postbreeding), but deliveries were preterm in Ovx mice within 24h (day 17). In the Ovx+P4 group, births were delayed to term and post-term in most Intact+P4 mice with adverse pregnancy outcomes commonplace. Characteristics of cell nuclei and degradation of cross-linked collagen were advanced with preterm birth in Ovx mice compared to controls that gave birth by at term. Treatment of Ovx mice with P4 blocked preterm birth, but parturition was complicated by dystocia. In addition, P4 given to ovary-intact mice sustained peak pregnancy concentrations, but had minimal effects on cytoarchitecture of the prepartum cervix stroma except term birth was forestalled with dystocia and fetal morbidity. Density of resident macrophages in the cervix stroma in term Ovx+P4 mice was reduced along with area of macrophage stain versus postpartum controls. Thus, analyses of cervix cellular cytoarchitecture provided useful biomarkers of local inflammation to assessment the ripening process for preterm and term parturition. Collectively, findings suggest a functional loss of prepartum cervix responses to progesterone are part of a final common mechanism for parturition across mammals.

**Summary:** Loss of response to progesterone withdrawal is associated with cervix ripening while some cytoarchitectural characteristics of remodeling are regulated to block preterm birth and dystocia at term

## Introduction

The landmark 1956 review by Csapo acknowledged the essential role of progesterone to sustain pregnancy and surmised that loss of response to progesterone was required for labor and birth at term ^1^. In women, some primates, and the horse, progesterone in maternal circulation remain elevated antepartum then decline after labor or postpartum ^2, 3^. In multiple mammalian models for parturition, progesterone in circulation falls occurs contemporaneously with the shift to the contractile uterine phenotype of labor ^4–8^. The essential importance of the protective effects progesterone to sustain pregnancy comes from clear findings that preterm birth follows either a major decline in maternal circulation, as with removal of the ovary (principal site of production in some mammals), or treatment with nuclear progesterone receptor antagonists ^9, 10, 23^. Limited availability of reproductive tissues in primates during the prepartum period leading up to labor at term or in subjects at risk for preterm birth present a significant challenge for investigations of the mechanism for parturition.

Accordingly, studies in rodents have provided useful insights about temporal changes in maternal circulation and reproductive tissues as pregnancy progresses to term. A precipitous fall in systemic progesterone is reported by the afternoon of the day preceding birth to coincide with labor in mice ^5, 8, 11, 12^. Before then, replicable evidence indicates that progesterone in maternal circulation is at or near peak concentrations of pregnancy ^8, 13 1, 14^. Plasma concentrations of multiple proinflammatory cytokines are at or below baseline throughout this prepartum period up until the day of delivery at term ^13^. Thus, evidence suggests an absence of systemic maternal immune system inflammation even with detectable presence of fetal allogeneic cells and molecules in circulation.

By contrast, the prepartum cervix cytoarchitecture appears to remodel though a process typically associated with local inflammation other tissues. Specific morphological changes in the cervix stroma 2-5 days before birth include a decline in cell nuclei density, degradation of cross-linked collagen, and increased presence of activated macrophages^23^. These features characterize the Phase 2 ripening that follows softening of Phase 1 cervix remodeling ^11^. A comparable study of cellular and structure in non-laboring women before preterm and term birth supports this conclusion ^15^. Though prepartum biopsies from primates are unavailable for studies of gonadal steroid regulation of cervix remodeling, in mice estradiol and progesterone capsules block ovariectomy-induced preterm birth ^16^. Progesterone receptor agonists also rescue pregnancy and mitigate inflammation by reducing degradation of cross-linked collagen, increase cell nuclei density, and reduced density of macrophages/cell nuclei in cervix stroma ^15–19^. Moreover, progesterone upregulates synthesis of collagen fibers in the extracellular cervix matrix ^57^ to sustains a softened minimally distensibility cervix structure ^8, 21^. These findings implicate the capability of progesterone to suppress local inflammation in the cervix during prepartum structural and cytoarchitectural remodeling to block preterm birth. However, removal of the progesterone capsule from mice on day 18 postbreeding that had been ovariectomized 2 days earlier and treated for with both estradiol and progesterone led to birth 24h later (day 19) without effect on cervix cytoarchitecture. Whether birth resulted from systemic withdrawal of progesterone in these mice with estradiol still present or an endogenous loss of responsiveness by the cervix stroma to this progesterone is not known.

To focus on the important role of progesterone in term and preterm birth, the first objective was to test the hypothesis that actual loss of progesterone advances the inflammatory drive of prepartum cervix ripening before preterm and term birth. Based upon past observations that prolonged treatment of pregnant mice without ovaries, the principal source of endogenous progesterone production, have increased fetal morbidity and dystocia at term ^20^, the second goal was to test the hypothesis that cytoarchitectural prepartum cervix remodeling required a decline in systemic progesterone. Evidence suggests that progesterone can forestall cervix remodeling to term both associate with loss of systemic progesterone -induced preterm birth and in intact mice given, but raise concerns about unintended consequences that impair cervix ripening, newborn well-being, and parturition.

## Methods

Adult nulliparous CD-1 mice were purchased from Charles River (Hollister, CA) and individually housed in the Loma Linda University vivarium in controlled conditions with12-h lights on at 7 am PST. All procedures were conducted under an Institutional Animal Care and Use Committee-approved protocol that conformed to NIH Office of Laboratory Animal Welfare guidelines.

### Role of progesterone in prepartum cervix remodeling for term and preterm birth

Mice were ovariectomized under isoflurane anesthesia on day 16 postbreeding (8-9 am PST) to remove the main source of progesterone and estradiol production that sustains systemic concentrations during pregnancy in this species ^16^. Before skin closure, a 1 cm Silastic capsule containing either vehicle or progesterone (Ovx or Ovx+P4, 0.1g in 0.1ml sesame oil; Sigma, St. Louis, MO) was inserted subcutaneously (s.c.) into the dorsal hindquarters as previously described ^21^. Sham-operated mice with intact ovaries and a sesame oil-filled capsule served as controls (Vehicle). Later that afternoon, designated day 16.5 postbreeding (about 4pm), prepartum mice in the Vehicle, Ovx, and Ovx+P4 groups (n=4-16, each) were asphyxiated with CO_2_. Thereafter, other mice in Sham, Ovx, and Ovx+P4 groups, respectively, were periodically checked for birth of at least 1 pup (postpartum PP). In addition, a group of day 15 postbreeding mice served as a pre-study baseline control.

To assess the effects of sustained progesterone on cervix remodeling at term, ovary-intact pregnant mice were anesthetized and implanted with a capsule that contained progesterone on day 16 postbreeding (Intact+P4, n=4-5/group). These mice were subjected to the same as protocol as described above, until pups were born or until day 22 when the study was concluded.

#### Sample processing

Postmortem, blood was collected and serum stored at -80°C for later progesterone assay (ELISA, Enzo Life Sciences kit #ADI-900-011 performed by KMI, Minneapolis, MN; sensitivity <8.57 pg/ml, intra- - and inter-assay coefficients of variation averaged <9%). From each mouse, the cervix with attached vaginal tissue surrounding the ectocervix, the endocervix, and a small portion of each uterine horn was excised and immersion fixed in fresh 4% buffered paraformaldehyde for 24h, then placed into 70% ethanol.

#### Image analyses of cross-linked collagen, cell counts, and macrophage morphometrics

Tissues were paraffin-embedded and sectioned in the coronal frontal plane (10 μm each) and. processed to assess cross-linked collagen using picrosirius red (Cat. No. 24901; Polysciences, Warrington, PA). As previously detailed, optical density of picrosirius red stain birefringence viewed with circular polarized light is specific and inversely related to degradation of collagen cross-linking ^16, 22, 23^. Data were normalized to cell nuclei/area to account for fibrillary structural remodeling of the cervix as expansion occurs with progresses of pregnancy to term. This analysis of collagen staining is the only *in situ* approach to specifically assess extracellular cross-linked collagen in the intact cervix stroma and, as mentioned in reviews, has proven useful in other fixed tissues, as well as supported by correlations with biochemical and electron microscopic analysis of fiber length and diameter in cervix stroma, as well as more recently with PA US^24^.

Adjacent sections were also processed by immunohistochemistry to identify dark brown DAB-stained F4/80- macrophages (1:800 dilution; T-2006; Bachem, Torrance, CA; (Cat #K3468; DAKO, Carpentaria, CA) with methyl green counterstained cell nucleus. Whole sections were scanned (Leica-Aperio CS2 scanner). A survey of 6-8 photomicrographs from the ectocervix and endocervix stroma of each mouse were captured (Imagescope software, Leica Biosystems, Deer Park IL) from 2 sections/mouse (12-16 snapshots in a total average area of 1.5-2×10^7^ µm^3^/mouse). In these regions of interest, lumen, mucus, epithelium, subepithelium, large blood vessels with cellular contents, and any atypical structures were excluded from analyses. Thus, in the cervix stroma, manual counts of cell nuclei and macrophages made using NIH Image J according to previously established rigorous criteria of morphological features that were independently replicated by multiple co-authors and research staff ^16, 21, 25, 26^. The coefficient of variation in replicate cell nuclei and macrophage counts/area by individuals was <10%. In addition, assessment of cell nuclei density in all ROIs were confirmed with an automated trained algorithm using Visiopharm software that was previously validated ^13^. In addition, scripts and algorithms for use in QuPath were developed to automate image analyses of area of F4/80 macrophage-specific stain. Validation of this approach was based up Brey et al^61^ and presented in the Supplement.

#### Statistical analyses

Macrophages/area and OD of collagen birefringence were normalized to cell nuclei density for each ROI to account for variations in cell sizes and extracellular space in cervix sections within individuals, as well as among mice in groups and with progression of pregnancy. Data were initially analyzed by Grubb’s and Levene’s tests to screen for outliers and variance of data. Data that were normally distributed (p>0.05 Levene’s test) were evaluated by one-way ANOVA with either Tukey’s test for individual post-hoc analysis or Dunnett’s test versus D15 Sham or D16.5 Sham-operated mice. Data not normally distributed were analyzed with Chi square and Kruskal-Wallace tests for individual comparisons. p < 0.05 was considered significant for all tests (PRISM 11; GraphPad, San Diego, CA).

## Results

### Progesterone blocks preterm birth with adverse pregnancy outcome with impaired parturition

Vehicle controls gave birth at term on average day 19.5 postbreeding by the morning of day 19 or 20 (Fig 1 Panel A). By comparison, removal of the ovaries from pregnant mice on the morning of day 16 led to birth within 24h on day 17 postbreeding, significantly advanced compared to term births in vehicle, intact mice and even later the Intact+P4 group. By contrast for progesterone-treated mice without ovaries, preterm birth was forestalled to a range of 18-20 days postbreeding. Effects of progesterone implants on parturition in intact mice (Intact+P4), resulted in an average day of birth of 20.5 days postbreeding, most before the morning, in 4 of 30 intact mice. Remaining mice in the Intact+P4 groups were pregnant until conclusion of the study on day 22 postbreeding.

**Figure 1.**
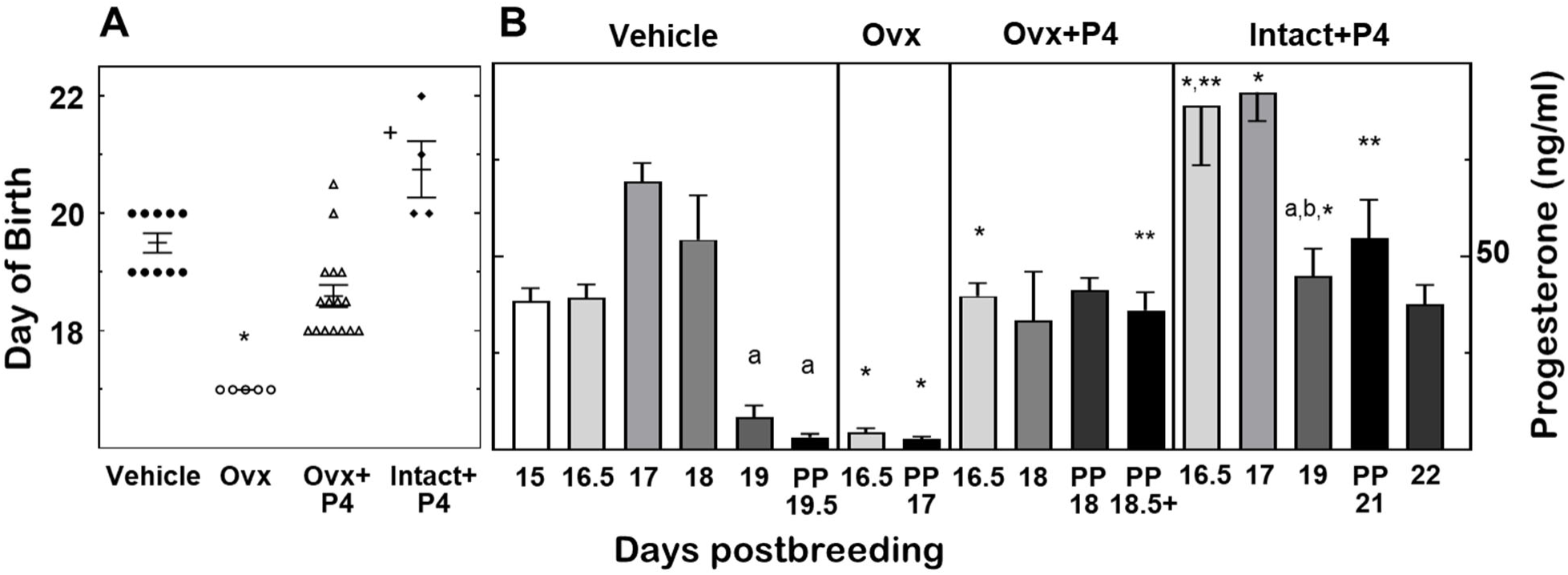
Panel A. Birth of at least 1 pups in Vehicle-treated intact controls (filled circles), as well as Ovariectomized mice given a capsule filled with oil (Ovx, open circles) or progesterone (Ovx+P4, open triangles), and Intact mice with progesterone capsule (Intact+P4, filled diamonds). Data are mean ±SE (average n=7/group, range 4-16). Statistical analyses included Grubb’s and Levene’s (p<0.05) tests which required non-parametric analyses. p<0.05 (Kruskall-Wallis with Dunn’s tests): *Ovx versus Vehicle, +Intact+P4 versus Ovx and Ovx+P4 groups. Panel B. Plasma progesterone concentrations in same groups of mice in Panel A. Data are mean±SE (average n=6/group/day, range 3-16/group/day) on various days postbreeding. p<0.05 (one-way ANOVA with Tukey’s test) within Vehicle controls versus a15-D18 and within Intact+P4 versus aIntact+P4 D16.5 and bD17. Comparisons among groups same day postbreeding Grubb’s and p>0.05 Levene’s tests were followed by parametric tests. p<0.05 (one-way ANOVA with Tukey’s test) *prepartum Ovx Day 16.5 and Intact+P4 D16.5 versus Vehicle D16.5 group; **both PP Ovx+P4 (PP preterm D18 and PP term D18.5+ groups), PP Day 21 Intact+P4, and delayed prepartum D22 Intact+P4 mice versus PP term Vehicle and preterm PP D17 Ovx mice.

Assessment of morbidity, mortality, and birth complications of pups and pregnant mice in vehicle-treated controls indicated an average litter size of 12 implantation sites or newborns across groups. In controls, there were an average of 12 pup/litters (6-14 range). All pups were pink, alive, had nursed (milk in stomach), and were present in the nest of bedding. Postmortem. 2 fetuses were reabsorbing at implantation sites in separate dams and one dam appeared lethargic postpartum. Compared to the 0.05% adverse pregnancy outcome in controls (3 of 60 mice), prepartum Ovx mice on day 16 postbreeding had a comparable compliment of fetuses in utero that were pink. By day 17, the 5 postpartum dams with PTB (PP Ovx) had an average of 12 implantation sites (6-15 range), with 12 of 61 fetuses intact, present in the cage along with cannibalized remains. Some preterm fetuses had a grey pallor and presumed to be stillborn while none had nursed.

By contrast, progesterone treatment of mice lacking ovaries increased incidence of adverse pregnancy outcomes. Prepartum, Ovx+P4 mice on days 16.5 and 18 postbreeding averaged 11 implantation sites/litter with comparable rates of resorptions in utero as that in controls and Ovx mice. However, in postpartum Ovx+P4 mice, of the 199 implantation sites in 16 dams, 78 pups were dead in utero, stillborn (grey) or missing in the cage and presumed cannibalized-a 32% adverse pregnancy outcome. Of the 103 pups that were born, all nursed though 9 exhibited some with contusions. In 2 postpartum dams that had already delivered the majority of pups, several fetuses remained undelivered in a contracted uterine horn with one pup lodged in the birth canal of each. These findings implicate dystocia in the parturition process in upwards of 50% of dams in this group. When the study concluded on the morning of day 22 postbreeding, 5 ovary-intact mice in the Intact+P4 group had not delivered. With an average of 12 implantation sites/mouse, all fetuses were in the process of resorbing in utero and 4 dams presented compacted uterine horns with a fetus lodged at the uterine funnel isthmus into the internal os. Maternal morbidity nor mortality was evident.

### Effects of ovariectomy or progesterone treatment on concentrations in circulation

Systemic progesterone in each of these groups met expectations of findings in previous studies with similar treatments ^5, 21, 27–29^. In prepartum Vehicle controls, progesterone was elevated in circulation on prepartum days 15-18 compared to that on prepartum day 19 and in postpartum mice (days 19-20 postbreeding) or in mice without ovaries given a vehicle-filled capsule (Fig 1 Panel B). Thus, progesterone treatment of ovariectomized mice sustained systemic concentrations of approximately 35 ng/ml, 6-10-fold higher than in controls both on the day before and day of birth, as well as in prepartum and postpartum ovariectomized mice with a vehicle implant. Mice in the Intact+P4 group had peak concentrations of progesterone in the circulation of vehicle controls occurred on day 17 of pregnancy. Systemic concentrations declined by the morning of prepartum day 19 to that found in postpartum mice, significantly reduced compared to earlier in pregnancy. By contrast, P4 treatment sustained peak concentrations. In these intact given P4, mice on days 16.5 and 17 postbreeding had increased progesterone compared to vehicle controls on the same day of pregnancy or day 15 of pregnancy in the experiment 1 while mice on days 19 and >21 postbreeding had increased concentrations relative to day 19 and PP Vehicle groups.

### Effects of ovariectomy and progesterone on cytoarchitecture and spatial morphometry of the cervix

Remodeling of cervix cytoarchitecture was evident while progesterone in circulation of pregnant controls was at or near peak concentrations (Fig 2). By prepartum day 17, density of cell nuclei declined and increased optical density of collagen birefringence increased in the prepartum cervix. By the day of birth, compared to prepartum days 15 and 16.5, the density of macrophages had also increased along with reduced cell nuclei and increased optical density. Each of these cytoarchitectural features reflect aspects of local inflammation in prepartum cervix remodeling.

**Figure 2.**
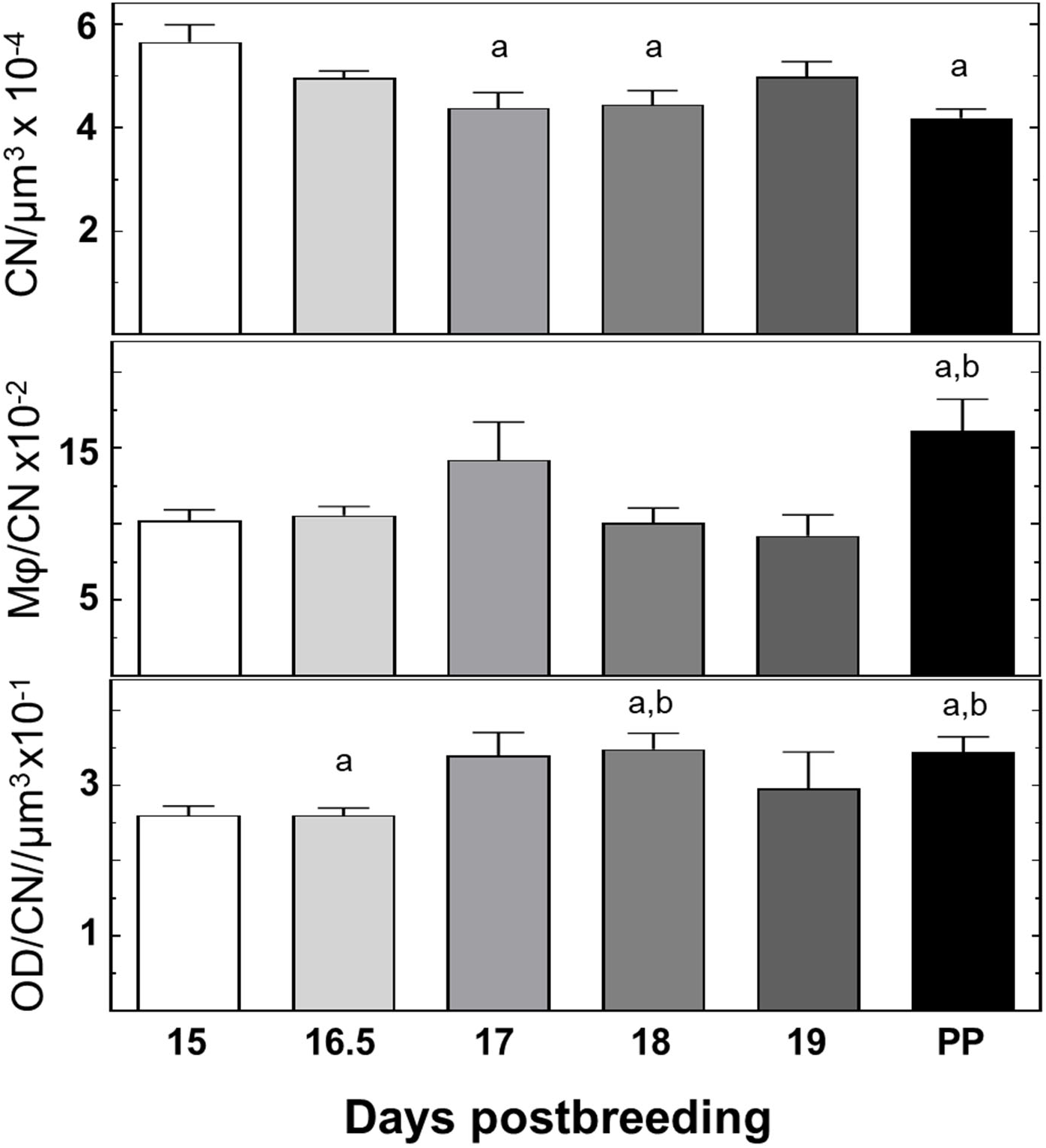
Prepartum declines in cell nuclei density and cross-linked collagen before term and in density of macrophage/cell nuclei by the day of birth indicate increased prepartum cervix inflammation. Data are mean ± SE (average n=10, range 4-16 mice)/group/day postbreeding) of densities for cell nuclei (CN/µm3), Mφ cell numbers/CNµm3, and cross-linked collagen (inverse of OD/CN/µm3). After Grubb’s and p>0.05 Levene’s tests for outliers and homogeneity of variance, respectively, to qualify for parametric statistics, data were analyzed by one-way ANOVA with Dunnett’s test across days postbreeding. p<0.05 versus aday 15 postbreeding (D15) or bday 16.5 postbreeding (D16.5).

In Ovx mice as in ovary-intact controls, cell nuclei density in the prepartum cervix was similarly reduced by day 16.5 postbreeding, but not in Ovx+P4 mice compared to that in day 15 controls (Fig 3 versus Fig 2). In contrast in Ovx+P4 mice by day 18, cell nuclei density increased in the cervix stroma compared to day 15 controls and Ovx day 16.5 mice. By term, cell nuclei density was declined in PP Ovx+P4 mice versus day 18 Ovx+P4 mice but still elevated compared to Ovx PP mice - a finding consistent with previous reports of reduced remodeling at birth.

**Figure 3.**
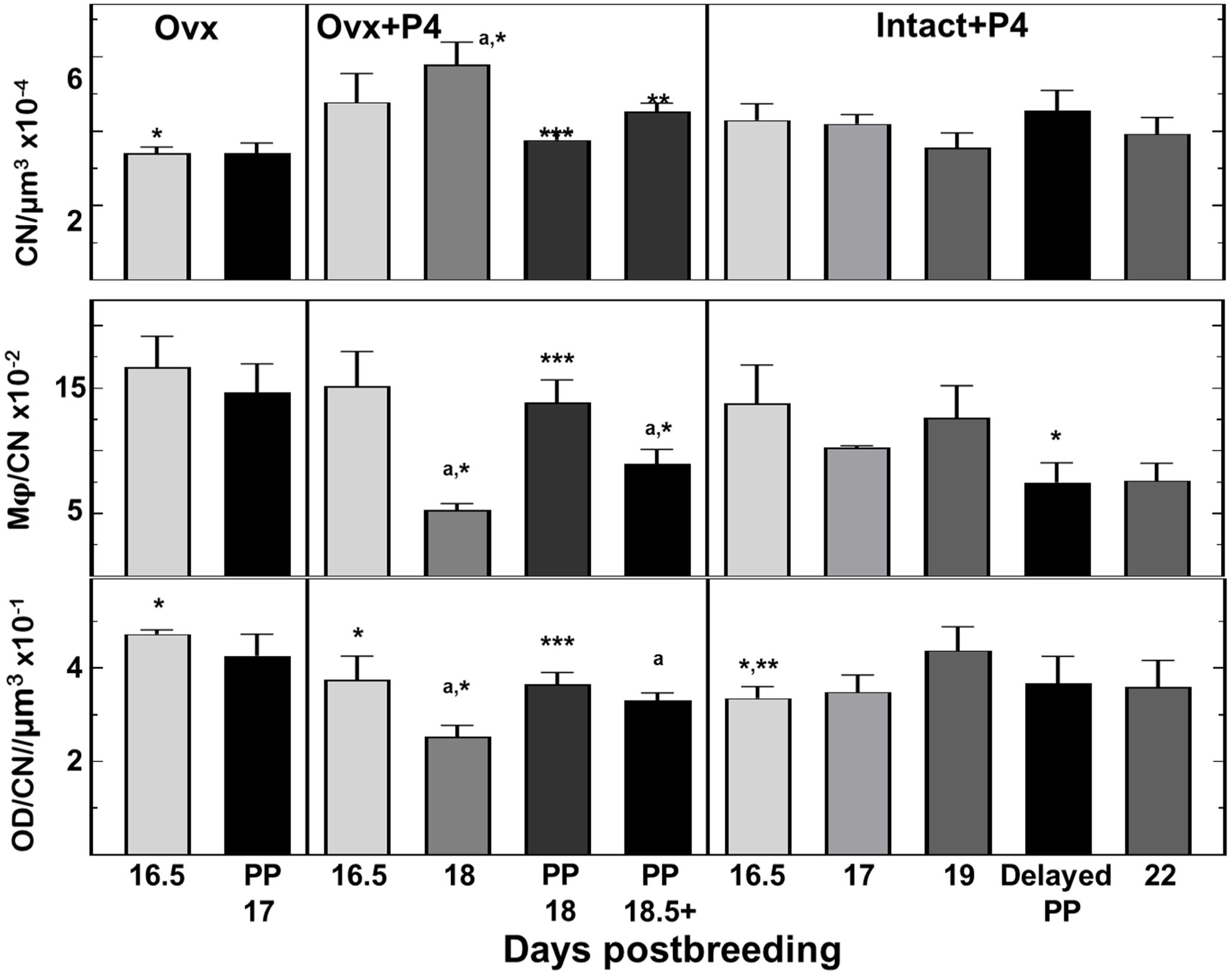
Progesterone sustains pregnancy in mice lacking ovaries, but impairs cervix remodeling and leads to adverse outcomes with birth at term. Quantitative assessment of cervix cytoarchitecture indicate decline in CN density and cross-linked collagen, but no change in density of Mφs with reduced P$ after ovary removal. Progesterone treatment after loss of endogenous production blocks these effects within 2 days on cell nuclei and cross-linked collagen and reduced density of Mφs in the cervix (Ovx+P4 D18 vs Ovx days16.5 and 17). However, responsiveness to P4 was diminished or lost and birth impaired at term. Groups and abbreviations are described in text and the legends of Figures 1 and 2. Data are the mean±SE (average of n=6, 4-9/group/day postbreeding). Mice were ovariectomized on day 16 postbreeding and given a vehicle-filled or P4-filled capsule (Ovx or Ovx+P4). Data passed Grubb’s and Levene’s tests for one-way ANOVA with Tukey’s analyses. Within each group across days postbreeding, p<0.05 indicated for ^a^CN/µm^3^, Mφ/CN, and OD/CN/µm^3^ for prepartum OVX+P4 D18 versus Ovx D16.5, as well as for ^b^Mφ/CN and OD/CN/µm^3^ in PP day 18.5+ mice versus Ovx day 16.5 and Ovx PP (D17) mice. Among groups on the same day postbreeding, p<0.05 indicated for prepartum CN/µm^3^ and OD/CN/µm^3^ data *D16.5 Ovx versus D16.5 controls, and for OD/CN/µm^3^ data for D16.5 Ovx+P4 and D16.5 Intact+P4 groups versus D16.5 Ovx, as well as prepartum CN/µm^3^, Mφ/CN, and OD/CN/µm^3^ data on D18 Ovx+P4 versus controls; for Mφ/CN delayed PP Intact+P4 versus PP term controls. For CN/µm^3^ ** Ovx+P4 PP D18.5+ versus Ovx PTB (D17) mice and D16.5 Intact+P4. For CN/µm^3^, Mφ/CN, and OD/CN/µm^3^ *** PP D18 Ovx+P4 vs D18 controls.

**Figure 4.**
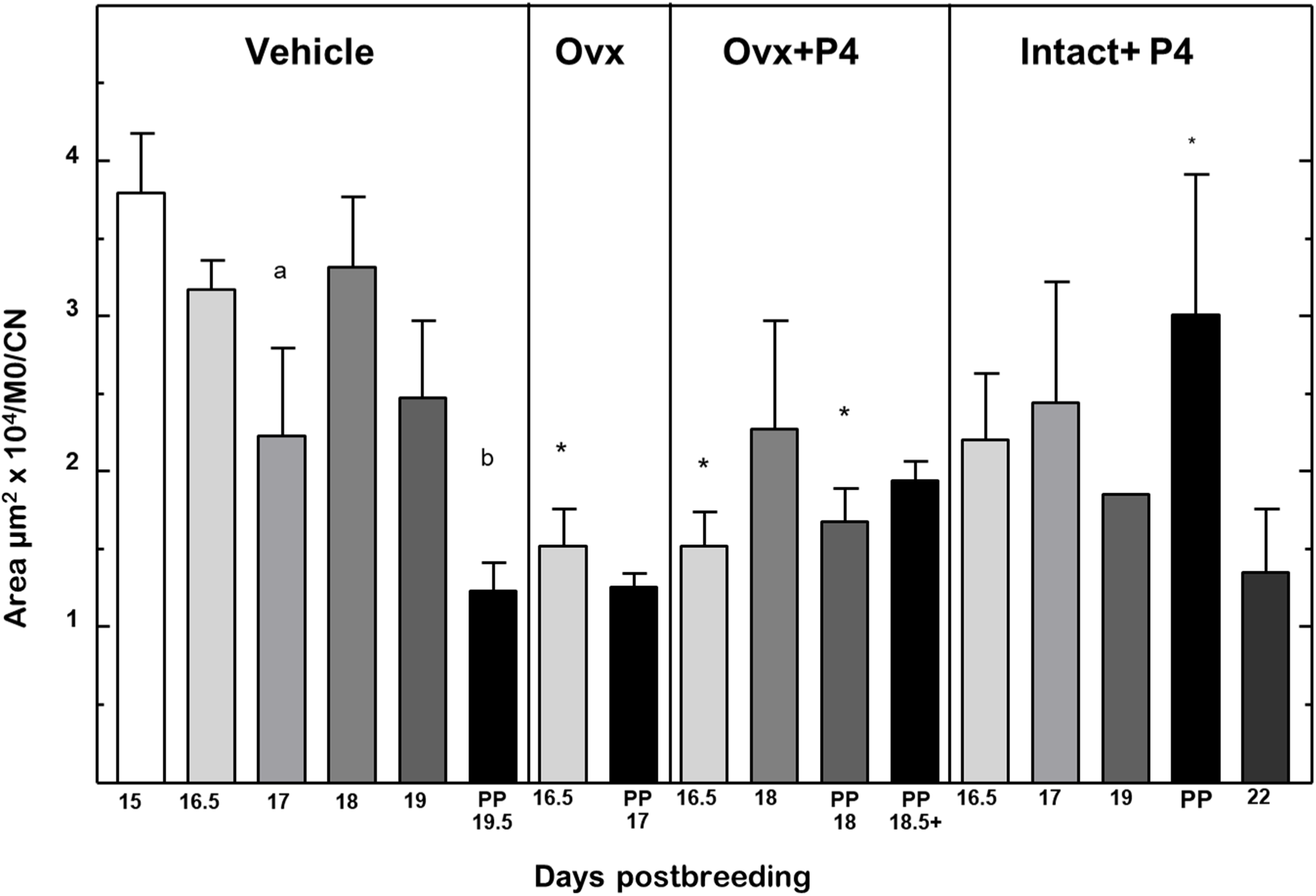
Area of macrophage stain normalized for macrophage cell number per cell nuclei density. Inflammation decline before birth at term and with PTB induced by loss of, but not regulated by progesterone. Groups are the same as described in Figure 1 legend. A script using QuPath software was customized to identify area of macrophage stain based upon color, intensity, and pixel numbers in a survey of photomicrographs in sections of the ectocervix and endocervix regions of interest (ROIs) from mice as described in Methods and validated in the Supplement. As an indication of inflammation related to the extent of presence of macrophages in the cervix, area of macrophage stain was normalized to number of macrophages per cell nuclei density within the ROI. Data are mean ±SE (n=3-15 mice/group/day, average 7/group/day), p<0.05 (one-way ANOVA. b versus D15, D16.5, and D18 within Vehicle group. Comparisons among groups same day postbreeding day * p<0.05 (one-way ANOV) verses Vehicle same day. For all other comparisons, data were not significantly different.

For macrophages, cell density increased in the cervix stroma of controls by postpartum vs day 16.5 groups. By contrast, in Ovx mice, cervix macrophage density declined within 6h of removal of ovaries and, remained low when compared to controls the next day for preterm birth on day D17. This suggests that PTB may not be required for PTB. Importantly, the decline in resident macrophages after removal of the ovaries was blocked by P4 treatment. Macrophage density in the cervix of Ovx+P4 mice was comparable to that in Vehicle controls and increased relative to the Ovx group on day 16.5 postbreeding.

For cross-linked collagen, the inverse of OD of picrosirius red birefringence, degradation increased within 6h in Ovx Day 16.5 mice compared to day 15 controls (Figure 3 bottom panel). However, in P4-treated Ovx mice, OD was not different in the D16.5 and D18 groups compared to each other or controls, but increased (less cross-linked collagen) by preterm PP day 18. Thus, intact and Ovx+P4 mice at term, as well as Ovx mice with preterm birth had increased OD birefringence relative to earlier on pregnancy - an indication that extracellular cross-linked collagen was reduced in the cervix stroma in time for birth.

With respect to the cervix morphology, cell nuclei density in the stroma of Intact+P4-treated mice, was unchanged throughout pregnancy and compared to that in controls. Macrophage density was also unchanged in the prepartum cervix stroma across all groups whether control or Ovx+P4 groups except reduced with delayed birth versus Ovx+P4 PP D18 (Figs 2 and 3 middle panels). No change in cross-linked collagen in prepartum cervix stroma was indicated for cross-linked collagen in the Inact+P4 groups after the decline in Itact+P4 mice on day 16.5 of pregnancy versus day 16,5 controls and the day 16.5 Ovx group. Thus, in prepartum mice, there was no evidence of response to P4 for the 3 characteristics of cervix stroma inflammation, cell nuclei density, cross-linked collagen, and macrophage density during the prepartum period leading up to term birth before day 20 of pregnancy.

## Discussion

The objectives of the present studies were to determine if withdrawal of progesterone efficacy advanced the inflammatory drive of cervix ripening for preterm and term birth. The first study eliminated the main source of progesterone production during pregnancy to reduce then restore with progesterone treatment physiological systemic concentrations. Removal of the ovaries induced preterm birth within 24 h while progesterone delayed preterm birth. These results replicate previous findings ^16^. In addition, progesterone delayed term birth in ovary-intact mice.

However, the birth process was substantially impaired for the majority of progesterone-treated mice in both studies as evidence of shortened length of uterine horns, fetal resorptions or compacted amniotic sacs, and a fetus was engaged with or in the lower uterine segment birth canal. Intrauterine hemorrhage was also observed in some post-term progesterone-treated mice, the likely result of difficulties with the birth process analogous to dystocia in women witht risk for fetal stress and adverse pregnancy outcome. These findings are consistent with a functional progesterone withdrawal by the uterus at term ^30, 31^. These findings also support a functional loss of response to progesterone by the cervix and suggest that the inflammatory drive for prepartum ripening may be independent of this hormone’s action when continuously present to term with a negative impact on neonatal viability.

In particular, functional withdrawal of progesterone to sustain an unripe cervix is supported by structural changes associated with ripening whether or not the hormone was present. This conclusion comes from the reduced cell nuclei density, increased optical density of birefringence (reduced cross-linked collagen), and increased presence of macrophages by term in both Day 20 Ovx+P4 and PP vehicle-treated intact mice versus the Day 15 group in the preterm study (Figure 3A-C, p<0.05). This lack of response to progesterone contrasts with effects of treatment on cervix by day 16.5. Thus, progesterone given early in the ripening process mitigated inflammation in the extracellular matrix of the stroma. Importantly, the capability of P4-treatment to forestall the decline in resident macrophages in the cervix in mice following Ovx replicates previous findings for effects of progesterone receptor agonists in Ovx mice ^16, 32^. Collectively, these data suggest that suppression of macrophage density and presumably inflammation are essential components for premature cervix ripening and preterm birth in Ovx mice. Moreover, as indicated in Figure 3B, progesterone sustained a resident macrophage population in Day 16.5 Ovx mice that contributed to the delay in preterm term. This indication is supported by a study that depleted and impaired macrophages in transgenic CD11b-*dtr* mice on day 15 postbreeding and found that macrophages are likely to be important not only for cervix ripening but to sustain pregnancy ^33^. These findings raise the possibility that progesterone treatments may regulate an alternatively activated macrophage phenotype that sustains an unripe cervix and forestalls prepartum preterm ripening.

In fact, comparison of gross and cellular features of the cervix across mammals indicate substantial similarities in the structure and function compared to diversity in other reproductive tract structures associated with pregnancy ^8, 15, 34, 36^. In particular, the cervix is distinct from the development, morphology, and function of the uterus ^23^, a fact concealed by common use of the term ‘uterine cervix’ ^35^. The predominance of connective tissue with an increasing gradient of smooth muscle from the ectocervix to uterine isthmus is dissimilar from the smooth muscle composition and structure of the uterus ^36, 60^. Further differentiating these two organs are their embryology, endocrine responses, resident immune cells, as well as degradation of dense fibrous cross-linked collagen with ripening, and enhanced innervation as pregnancy nears term^23^. Questions about structural remodeling of the cervix as distinct from the uterus are important since a carefully designed study by the Olson’s group indicated that prolonged labor and delayed delivery of the first pup past term occurred when concentrations of progesterone in circulation of ovary intact dams were sustained^5^.

Results of the study of Intact +P4 treated mice also support the conclusion that inflammatory processes in the extracellular stroma reflect a functional loss of response to progesterone at term. Each of the characteristics associated with prepartum ripening in ovary-intact mice were not affected by P4 treatment (Figures 5 A-C). At term, comparison of endpoints in PP intact controls with mice on day 15 postbreeding from the first study indicate ripening had occurred (fewer cell nuclei, increased resident macrophages, and less cross-linked collagen per area (Figure 5 vs 3, PP vs day 15 groups). However, compared to day 15 controls, in the Ovx+P4 group on day 20 postbreeding, neither cell nuclei nor cross-linked collagen were altered by P4 treatment in the cervix from >D21 mice; only macrophage density was reduced. These findings raise the possibility that progesterone regulation of cervix structure and either macrophage density or phenotypic activity at term may depend upon other ovarian products. Although beyond the scope of the present study, the idea that proinflammatory molecules from leukocytes may mediate effects of progesterone withdrawal for cervix ripening in the rat has been previously suggested ^18^. In a broader context, regulation of the onset and duration for loss of response to progesterone by the cervix must take into consideration the role of progesterone receptor A isoform, which is predominantly expressed by stromal fibroblasts, not macrophages ^25, 26^,, and presumably regulated by multiple factors, both endocrine and neural ^37^, before labor and birth in the mechanism for parturition.

Support for a functional progesterone withdrawal in both the preterm and term studies in this report is complicated by the high incidence of what appeared as dystocia and perinatal mortality in progesterone-treated mice with or without ovaries. Specifically, increased density of cell nuclei, fewer macrophages, and reduced cross-linked collagen in the cervix from P4 treated intact post-term mice on days 21 and 22, as well as in vehicle-treated PP mice compared to the day 15 controls (Figure 5). Lack of an increase in stromal macrophages with P4 treatment suggests an impaired remodeling process by the conclusion of pregnancy. After ripening, it is conceivable that responsiveness to progesterone may have been restored during the 3^rd^ phase of remodeling when dilation and effacement occurs with labor before preterm birth or at term. In Ovx and controls at term, systemic progesterone was reduced to baseline, so responsiveness to progesterone would not be a consideration for parturition preterm or at term, respectively. However with sustained progesterone in circulation from P4 treatment, dystocia occurred-a finding consistent with the ability of medroxyprogesterone, a progesterone receptor agonist, given with estradiol, to forestall ovariectomy-induced preterm birth, as well as block term birth in mice ^38^.

Prolonged pregnancy and complications in parturition associated with progestational treatments may solely affect local inflammatory processes that suppress cervix ripening. However, the possibility that progesterone affects the mechanism for maternal myometrial activity of labor, or act on the fetus or regulation of steroid metabolism in fetal membranes or decidua cannot be excluded ^39, 40^. Dystocia and lengthened pregnancy may also be an indirect consequence of progestogens promoting anti-inflammatory macrophage phenotypes or reported effects on other immune cells in the reproductive tract to disrupt cervix ripening ^41^. Whether identification of biomarkers for dilation and effacement, the third phase of prepartum remodeling, may help identify women at risk for preterm birth that may benefit from progestational treatments ^42^ or are in need of progesterone receptor antagonists to facilitate birth at term ^43–45^ to safeguard maternal and neonatal well-being remains to be determined.

The translational relevance of animal models is an important consideration to understand the mechanism for parturition in humans. First, there are more similarities than differences in the structure and function of the cervix among mammals as a barrier for protection of the developing fetus and its eventual elimination for birth to occur ^23^. Yet even among rodents, similarities in the Phase 2 ripening process between mice and rats ^15, 16^ during peak progesterone in circulation may not necessarily extend to dilation (Phase 3) or labor. The dystocia related to sustained systemic progesterone at term in the present study may reflect the importance of the decline in ovarian production of this hormone^27^, while in rats, both the ovary and placenta have this capability and placenta hypertrophy can sustain pregnancy following ovariectomy^7^.

In women, unique biomolecular markers have yet to be identified to reliably predict the progress of cervix remodeling ^46, 47^ or identify individuals at risk for premature or forestalled ripening have not been included in the current standard of clinical practice ^48^. Yet for some time, a modified version of the Bishop score has been repurposed to define cervix ripening. Though based upon empirical evidence of vaginal delivery after induction of labor in multigravida women at term, current use is untethered to structural, biomolecular, or physiological characteristics associated with ripening, which from the present consensus in animal models occur well before term. Availability of prepartum cervix before preterm or term is a limitation, though advances in non-invasive imagine technology that are validated to assess changes in cross-linked collagen, cell nuclei density, and inflammatory processes may prove useful to evaluate progress in cervix remodeling in pregnant women at term or as an endpoint for the effectiveness of treatments to reduce the incidence in women at risk preterm birth ^49–52^. The rationale for this approach is founded in present findings that constitute the first evidence for loss of response to progesterone by the cervix as reflected by characteristics associated with the inflammatory drive of ripening before term. Conceivably, increasing incidence of cesarean deliveries following induction of labor may reflect problems with sustained systemic progesterone and impaired cervix remodeling^59^. Thus, the present study supports the conclusion that loss of the trophic effects of progesterone could represent a common early sign for activation of the mechanism for parturition well in advance of changes in placental function ^53^, fetal membrane integrity ^54^, or increased contractility by the uterine myometrium during early labor ^55, 56^.

## Acknowledgments

Images were acquired for analyses in the California Tumor Tissue Registry Center in the Department of Pathology and Human Anatomy and the Advanced Imaging and Microscopy Core at the Loma Linda University School of Medicine. This project was in part supported by NIH HD054931, the LLU Department of Pediatrics Research Fund, and NSF MRI-DBI 0923559. The efforts by Anne C. Heuerman for help with data acquisition and analyses, as well as portions of each study are appreciated.

## Author’s roles and agreed order of list

**Steven M. Yellon** conceived, designed, and supervised experiments, as well as wrote the manuscript

**Daylan Ward** co-author performed data analyses, supervised statistics and graphics, co-wrote Results and parts of Discussion.

**Ashley Thompson** co-author data management, implemented statistics and QuPath data analyses, composed graphs and wrote figure legends, and referenced document

**Brigitte Vazquez** co-author contributed to the design and implementation of the experiment, performed statistical analyses, created data graphs, and edited the manuscript.

**D. Daniel Baldwin** developed algorithms and script for use and validation of QuPath image analyses of macrophage stain data, co-wrote findings and abstract

**Elaine J. Oldford** Co-author helped to do first study, worked up data, wrote initial draft

**Michael A Kirby** shared supervision and participated in conduct of experiments, data and manuscript review, contributed resources to fund study

---

**Supplement Figure 1.**
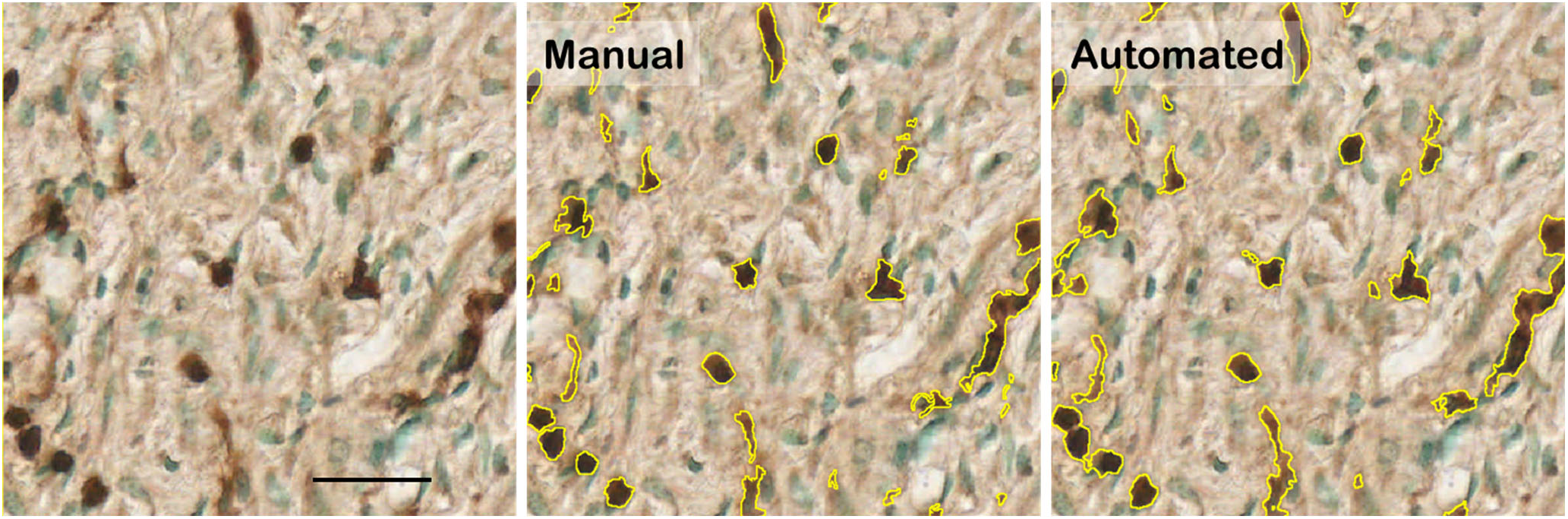
Example of comparison of original, as well as annotated Manual and Automated script (pODT) annotated area of macrophage stain in same photomicrograph of cervix stroma region. Scale bar =50µm

**Supplemental Table 1.** Automated approached to quantify area of macrophage-specific stain in immunohistochemical processed sections as previous detailed^61^. Comparison of Manual and Automated annotations of stain area (average of 3 trained Individuals assessing 35 photomicrographs in regions of cervix each). Correlational coefficient (*ρ*), average difference, bias correction factor (C_B_), and 95% confidence intervals were determined with a scripted Processed Optical Density Thresholder (pODT), a trained Pixel Classifier algorithm (PC), a scripted Optical Density Thresholder (ODT), and a trained Processed Pixel Classifier algorithm (pPC). *p*-values are Students t-test, which assessed the paired difference for masks from Manual and Automated approaches that quantified macrophage stain area after passing Levene’s test for normalcy of variance (p>0.05, only ODT data were log-transformed).

| Approach | Concordance<br>Correlational<br>Coefficient ($\rho$) | Average<br>Difference<br>$\mu\text{m}^2$ | Bias<br>Correction<br>Factor ($C_B$) | 95%<br>Confidence<br>Interval | $p$-value |
| --- | --- | --- | --- | --- | --- |
| pODT | 0.892 | 1.80 | 1.000 | 0.798 – 0.944 | 0.930 |
| PC | 0.697 | 72.29 | 0.950 | 0.491 – 0.829 | 0.024 |
| ODT | 0.613 | -92.80 | 0.907 | 0.391 – 0.767 | <0.001 |
| pPC | 0.411 | 257.47 | 0.531 | 0.247 – 0.552 | <0.001 |

**Supplemental Figure 2.**
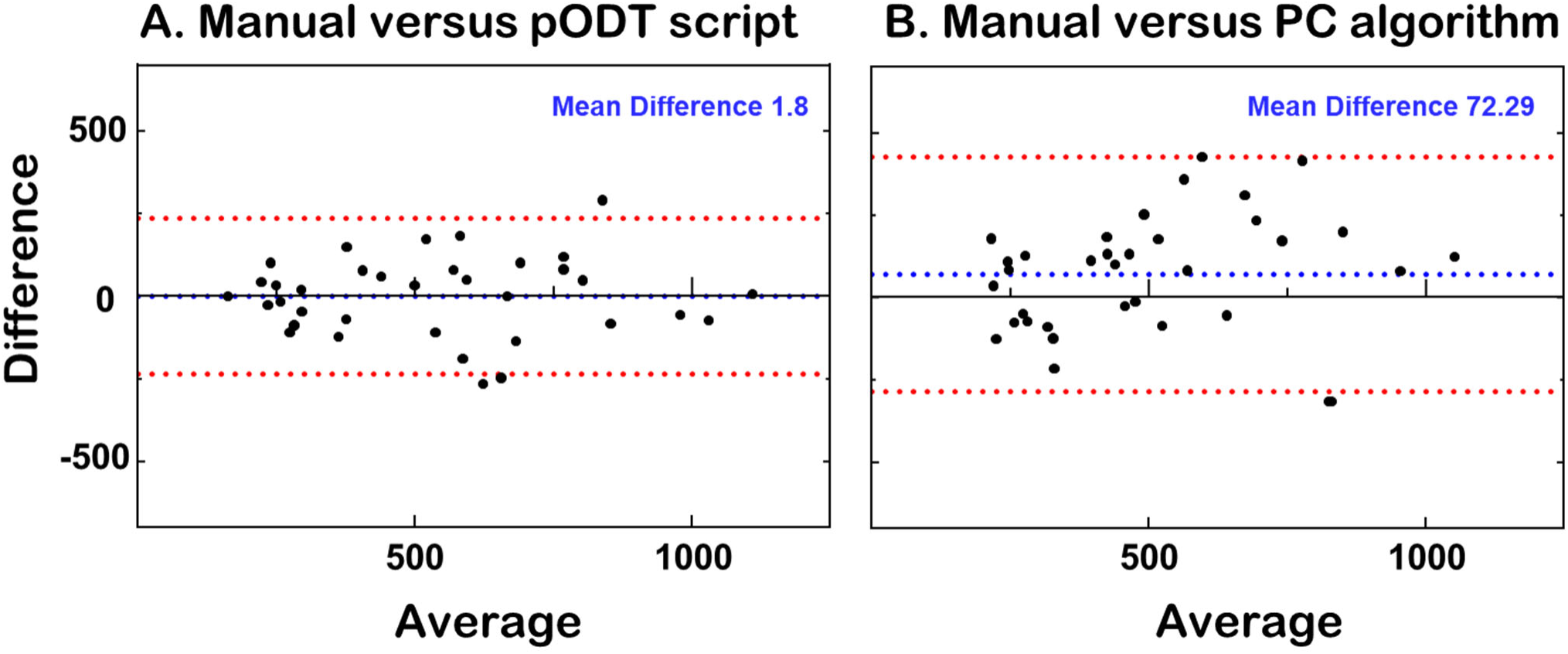
Bland-Altman plot analyses of differences in area measurements between the Manual annotated masks of 2 independent individuals and automated analyses with QuPath busing the pODT script model (Panel A) and the PC algorithm model with artificial intelligence and machine learning (Panel B). See Brey et al for details of validation of methods^61^.

